# Archaeological Ivories: A practical guide for identifying elephant and hippo ivory in the archaeological record

**DOI:** 10.1101/2024.01.27.576203

**Authors:** Harel Shochat

**Affiliations:** School of Archaeology and Maritime Cultures, University of Haifa, Haifa 3498838, Israel; Zinman Institute of Archaeology, University of Haifa, Haifa 3498838, Israel

## Abstract

The use of osseous materials for crafting various artifacts is well-attested in the archaeological record of the ancient Near East. While bone, the most common and available material, was used more frequently, ivory was highly valued as an exotic material for fashioning exquisite prestige items designed for ostentatious display and linked to society’s elite strata. Therefore, reliably distinguishing between the taxonomic origins of the osseous artifacts is significant for interpreting the archaeological data. In the Near East, the demand for ivory products reached its zenith during the mid-2^nd^ to mid-1^st^ millennia BCE, producing one of the largest assemblages unearthed anywhere in the world. Using state-of-the-art bio-molecular analyses to study this collection is costly, time-consuming, and entails damaging the highly-curated artifacts. This paper provides an essential, simple, non-destructive, and validated method to distinguish bone from elephant or hippo ivory (the most common osseous materials attested in the Near East record). However, following similar guidelines can help index other biological sources of ivory.

## Introduction: the study of ivory in the archaeological record

Ivory was an integral element in the Southern Levant material culture from the mid-2^nd^ to the mid-1^st^ millennium BCE (ca. 1600–600 BCE). The local demand for ivory artifacts reflects a broader phenomenon that characterized contemporary Near Eastern societies (Barnett 1982). The color and texture of the tusks of African and Asian elephant and hippopotamus teeth, their rarity, and their amenability for delicate carving anchored these materials’ prestigious status and symbolic prominence. Ivory became a charismatic medium for displaying power and authority probably due to its exotic value, qualities as a raw material suitable for various uses, or because it embodied human superiority over the largest terrestrial mammals.

Previous scholarship addressed the high cultural value of the material, assessing its social role and inclusion in the prestige economy mainly via interpretation of the artifacts’ typology, style, and artistic traits (for example, Mallowan and Davies 1970; Barnett and Davies 1975; Barnett 1982; Herrmann 1992; Feldman 2006, 2014). Yet, the potential embedded in the raw material analysis has been primarily understudied. As it was nonindigenous to the Southern Levant, in the case of elephants, and only possibly local in the case of hippos, identifying the artifacts’ raw material is essential for studying exchange mechanisms, reconstructing social stratification, and investigating biological and ecological aspects of ivory exploitation (for example, the question of the existence of the ‘Syrian’ elephant as a distinct sub-species; Collon 1977; Çakırlar and Ikram 2016). One of the paradigms in Near Eastern ivory research establishes the beginning of the ivory trade during the Late Bronze Age (*LBA*, ca. mid-16^th^ to mid-12^th^ centuries BCE) as a local exchange phenomenon (Caubet 2013) since hippos are said to be extant in the Levant up to the 10^th^ century BCE (Kolska-Horwitz and Tchernov 1990); thus ivory is theorized as being exchanged through local supply chain networks. Only later, during the Iron Age (*IA*; 11^th^ to early 6^th^ century BCE), when hippos became extinct, did exchange networks shift and expand to include ivory of elephant origin to meet the growing demands. Taxonomic identification greatly assists our understanding of exchange mechanisms and supply chain networks, whether local, inter-regional, or any hybrid mode, and directly impacts the reconstruction of past societies’ socioeconomic structures and intercultural spheres of interactions.

### Bio-molecular analysis of ivory

While traditionally, the word ‘ivory’ denotes the tusks of elephants, anatomically, these do not differ from the large teeth of any mammal, whether hippopotamus, wild boar, walrus, or others (Baker et al. 2020). As with any tooth, its primary component is dentine, a hard, calcified tissue composed of bioapatite crystals deposited in an organic matrix of collagen proteins (the main structural protein in the mammalian body; Hillson 2005: 184–193). Collagen’s durability and its relatively high degree of preservation, even when buried for millennia underground (similar to the almost entirely mineralized enamel tissue; Kendall et al. 2018; Weiner 2010), make it a potential candidate for bio-molecular investigations, such as proteomic and stable isotopes analyses.

The proteomic analysis or ZooMS (Zooarchaeology by Mass Spectrometry) method has been used in archaeology for over a decade now and has successfully provided taxonomic identification for faunal remains (mainly bones) at the family and genus levels (Buckley et al. 2009, Buckley 2018). It specifically targets those bones whose distinguishing morphological features were removed (for any reason, such as due to carving or crushing), thus making them unidentifiable. For that reason, this method is also relevant for identifying ivory artifacts.

The initial undertakings of stable isotopes for investigating archaeological ivories potential sourcing pools (for example, Nocete et al. 2013 study on earlier ivories of the third millennium BCE discovered in the Iberian Peninsula; African ivory exchange spheres from the 7^th^ to 10^th^ centuries CE by Coutu et al. 2016b; the East Africa ivory trade during the earlier modern era by Coutu et al. 2016a; and the west African trading routes by de Flamingh et al., 2021) has followed the footsteps of stable isotopic applications for modern ivories (for example, Vogel et al. 1990; van der Merwe et al. 1991; Koch et al. 1995; Cerling et al. 1999; Cerling et al. 2008). The investigations of modern ivories initially aimed to study the ecology and behavior of elephants and hippos in nature and later became an established forensic means to fight the illegal ivory trafficking (Ziegler et al. 2016).

Taking a similar forensic path, the Southern Levantine Ivory Research Project (LIRA) aimed to reconstruct inter-regional exchange networks of the mid-2^nd^ to mid-1^st^ millennia BCE, inter alia, by applying stable isotopes and ZooMS analyses as geospatial tracers for the raw material origin. The targeted database included thousands of potential artifacts, encompassing a chrono-stratigraphic sequence of a millennium of dense settlement history and spatially distributed at 46 sites encompassing all the geo-cultural units of the Southern Levant (Shochat 2023).

Considering such a broad scope of ivory and bone artifacts, the relatively high cost and resource consumption of the combined bio-molecular analyses necessitate a prescreening process. Seeking a practical prescreening method was further motivated by a tendency of scholars to unjustifiably label osseous artifacts as ‘ivory’ only due to their artistic quality (and in contrast, quotidian artifacts were assumed to be made of the ‘inferior’ bone; a repeated mistake in published archaeological reports and unpublished inventory lists).

This publication can partly be viewed as an attempt to create an essential tool for identifying ivory micro-structure using a straightforward method employable by any scholar. Practicality is the main objective of the method applied, and it is designed to be used anywhere, in the field or the lab, requiring minimum available technological and financial resources. It comprises two observational levels: macroscopic, using the naked eye, and microscopic, using a simple microscope (such as a portable handheld digital microscope). This method attempts to be straightforward and has a proven success rate, as substantiated by the proteomic-ZooMS analysis.

### Previous research on ivory micro-structure

The majority of the literature on ivory micro-morphology, mainly of elephant tusks, aims to explain the formation and structure of ivory or dentine tissue at its particle level (for example, using Scanning Electron Microscopy, SEM; Raubeheimer 1999; Su and Cui 1999; Virág 2012; Albéric et al. 2017; and most recently, Herron 2022). This direction of study is usually not applicable in archaeological research, as the required analytical equipment is expensive, thus unavailable in most archaeological labs, and requires technical training.

A small number of identification guides have been published. Two comprise the foundation course for the methodology used here, as they are both practical and simplified. Although elaborate and instructive, both suffer from some deficiencies for the abovementioned purpose.

The first is the Convention on International Trade in Endangered Species of Wild Fauna and Flora (*CITES*) and World Wildlife Fund (*WWF*) *Identification guide for ivory and its substitutes*, which was initially published by Espinoza and Mann (1992) and recently updated by Baker et al. (2020), is now accessible online, offering illustrations and sketches in color. This guide is directed to identifying modern ivory seized during the struggle against the commercial ivory trade. Targeting modern material, it focuses on identification features that are mostly unattainable when examining archaeological ivories owing to their irregular taphonomic history. For example, the essential identification step is to examine the suspected ivory object under long-wave UV light, as only genuine ivory will fluoresce brightly (Baker et al. 2020: 7–8, figs. 1.2A, B). Nevertheless, complete or broken, the surface of archaeological artifacts that have been buried underground for thousands of years does not necessarily glow under UV light.

In most cases, the surface of such artifacts is not polished, as they might have been on carving, if not treated or covered (colored, inlaid, or gilded). As a result, we often cannot observe some of the taxonomically indicative structural patterns we seek (such as the Scherger pattern, see below, and compare with Baker et al. 2020: figs. 2.6, 2.7). Moreover, in the archaeological record, we only occasionally find ivory in its original form – complete or large fragments of tusks or teeth. Discriminating taxa using a complete transverse cross-section of the tusk or tooth as a primary feature is thus less effective (see Baker et al. 2020: figs. 2.1, 6.3, 6.4, 7.3). One must adopt a three-dimensional perspective to correlate artifacts’ facets with the original tusk or tooth transverse (Tr), tangential (Ta), and radial (Ra) profiles (see Fig. 1).

**Fig. 1.**
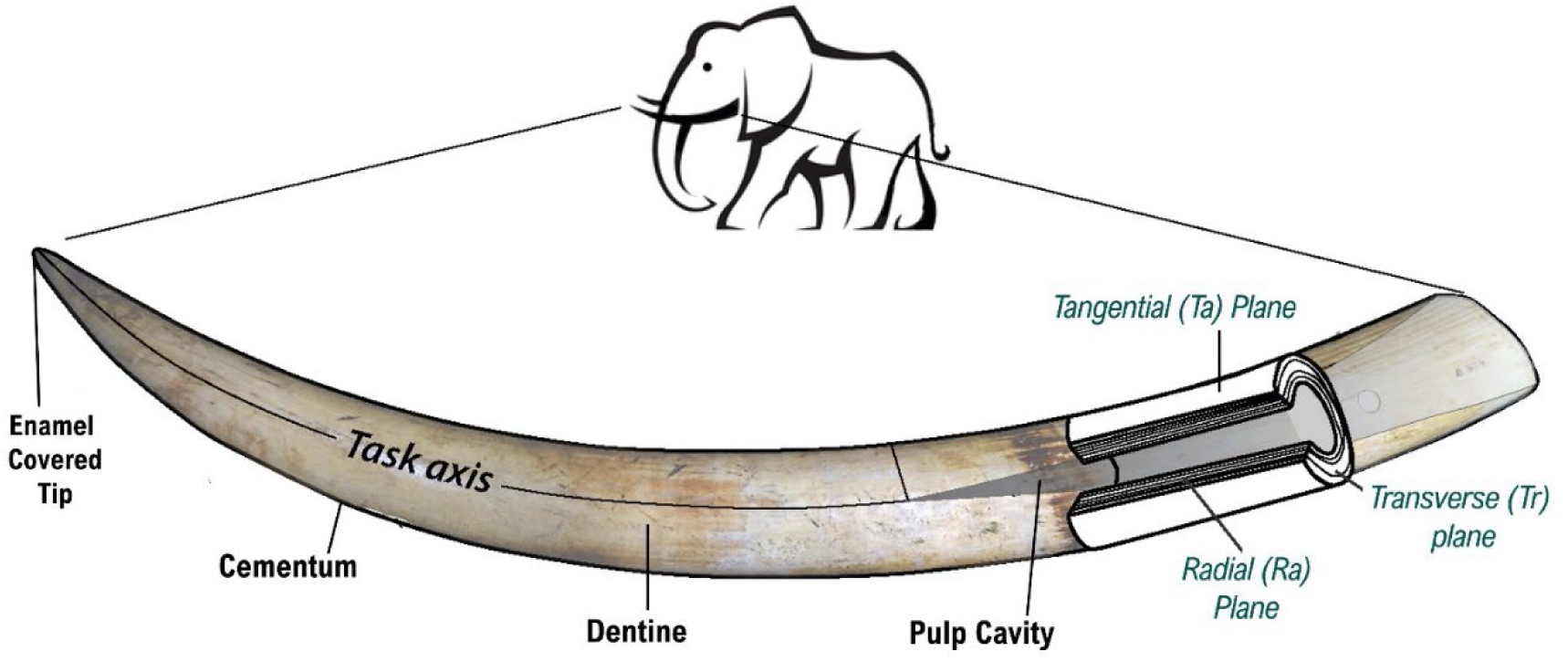
Basic structure and terminology of elephant tusk. Transverse (Tr), tangential (Ta), and radial (Ra) planes refer to the longitudinal tusk axis, and the breakage patterns along these plains are distinctive for each ivory, thus enabling the taxonomic identification of the raw material (illustration by Sveta Matskevich). Notice that the enamel layer covers only the tusk tip and is usually worn off by adulthood.

The second guide, an essential guidebook for dealing with ancient ivories in the Near East, is Krzyszkowska’s (1990) *Illustrated guide to ivory and related materials*. This guide, oriented toward archaeological practice, and despite its textual quality, it is visually deficient, considering it is over 30 years old.

Apart from these two manuals, the current identification method relies significantly on Locke’s (2008) seminal publication, probably the most thorough investigation to date, on the structure of ivory from the micrometer to the centimeter level. The lower resolution observations (at mm and cm scales) facilitate a three-dimensional outlook, enabling the study of the micro-morphological characteristics discernible in modern ivory and most ancient artifacts.

## Results: A straightforward and systematic ivory identification

The results described here are based on observations undertaken while examining no less than 1600 osseous artifacts (Shochat 2023). The notes taken through the initial stages of microscopic examinations were then converted into a three-stage protocol, as described below.

The microscopic observations were then compared to ZooMS analysis conducted on selected ivory and bone artifacts (N=88), aiming to evaluate the credibility of the microscopic method, out of which collagen was extracted successfully for 66% of the artifacts (N=58).

### Terminology and orientation for inspecting an ivory artifact

Teeth and tusks are physiologically the same tissue, differing in their growth mechanisms and natural function (Sukumar 2003). Their physical structure comprises a pulp cavity, dentine, cementum, and enamel (Fig. 1).

The pulp cavity, the innermost part, contains organic soft tissues (such as the nervous system) and is lined with odontoblastic cells responsible for dentine production. The organic (collagen fibrils) and inorganic (bioapatite crystals) materials composing the dentine are permeated along micro-canals, known as the dentinal tubules, which radiate from the central pulp outwards, extending to the dentine periphery, and its junction with the exterior cementum layer (Baker et al. 2020: 4–6). The formation and organization of these microscopic tubules (their elliptical profile major axis diameter is ∼ 3.5 μm) is a key element in the configuration of the hierarchical dentine structure (Raubenheimer et al. 1998: figs. 4–8; Raubenheimer 1999; Virág et al. 2012: figs. 4–6). In fact, with a simple microscope and magnification up to X 60–70 employed here, they cannot be viewed directly. However, the tubules alignment, stacked into microlaminae (micro-thin plates, sheets, or planes), their shape, orientation, and positioning in relation to the tusk axis (Fig. 2) are characteristic for each kind (taxon) of ivory (Locke 2008: 425–6, fig. 4). Consequently, they create distinctive and unique micro-morphological patterns that can be seen on the carved artifact’s surfaces (’under’ modern carving marks) or along its breakage planes, and constitute the principle means of differentiating elephant from hippo ivory.

**Fig. 2.**
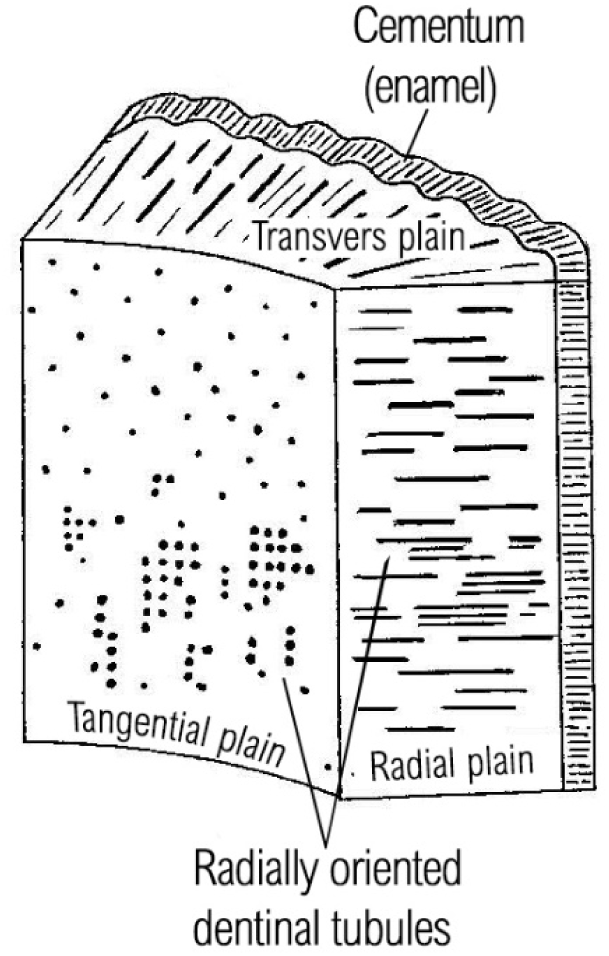
Schematic view of the radially oriented microlaminae formation (adapted from Locke 2008: fig. 7)

Another instrumental feature is the annual growth bands, which are observed as concentric rings in elephant tusks or hippo incisors but also adopt a triangular shape in hippo canines (see below). The long bone may also display the same concentric pattern, so these might be less indicative of our purpose. Nevertheless, the interface between the concentric bands and the perpendicularly oriented microlaminae creates zones of mechanical weakness due to cracking, creating distinct breakage patterns (Heckel 2018: 309–10). Breakage along the concentric bands also exposes the tangential plane of the incrementally growing teeth.

The upshot is that some variables are at play when inspecting ivory artifacts for their taxonomic origin (general bone, elephant tusks, or hippo canines and incisors). The first is that ivory artifacts were carved at an unknown angle to the general tusk axis (for suggested reconstructions according to previous scholarship, see Caubet and Poplin 1987: figs. 8, 9; Krzyszkowska 1990: fig. 28; Caubet and Gaborit-Chopin 2004: figs. 2–4). Therefore, we must first establish the correlation between the artifact’s facets and the original tusk or tooth transverse, tangential, and radial profiles (relating to the longitudinal axis) by recognizing the characteristic micro-morphological patterns typical to each taxon on the artifact’s surface or along breakage planes. The second variable is the diagenetic effect. The ivory artifacts have been deposited underground for thousands of years after finishing their life cycles and subjected to biological and physical degradation that might have altered or concealed the patterns we seek (for elaboration on the as-yet undetermined operating diagenetic mechanisms, see Kendall et al. 2018).

*Distinguishing bone from elephant and hippo ivory in three steps^1^*

**Step 1**. The macroscopic level: evaluating the artifact’s size, shape, surface features, and breakage patterns using the naked eye.

***Step 1: 1*** Observing the artifact’s general shape

The general shape of the larger ivory objects might reflect the original morphology of the teeth or tusks. An example is the ivory clapper found in Shiqmona, carved from a hippo lower canine (Fig. 3: A–B). Sometimes, although the raw material was reshaped, it is large enough to indicate its taxonomic origins, such as the furniture parts discovered at LBA Hazor (Fig. 3: C–D). Their sizeable dimensions indicate that these could only have been carved from elephant tusks (as for other indicative morphological features, see below).

**Fig. 3.**
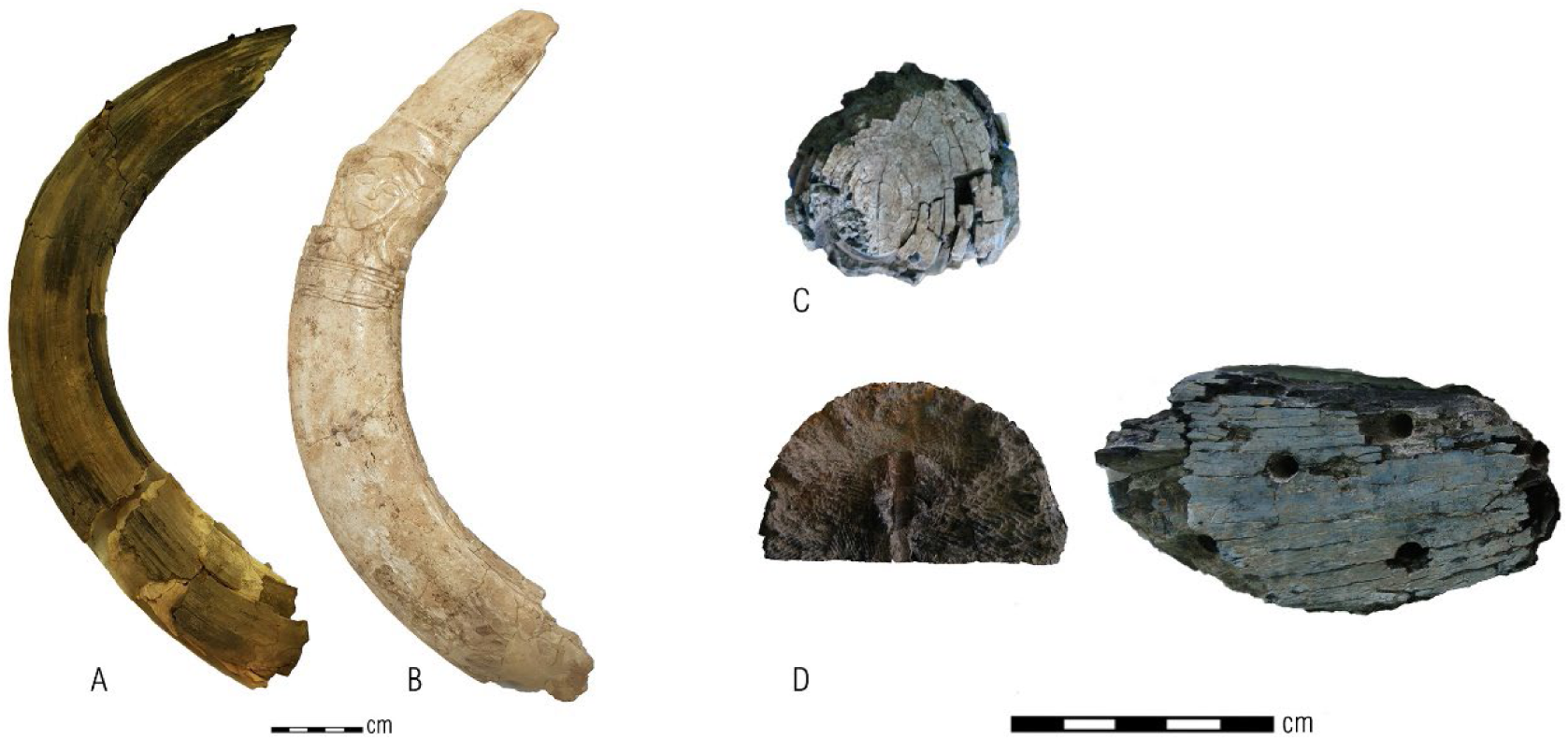
Ivory artifacts that retain the raw material general morphology. **A)** Unworked complete hippo lower canine found at Hazor (Hzr.81530); **B)** Ivory clapper (a cultic object) found at Shiqmona (Shq.11350); **C)** Large pieces of elephant tusks carved as furniture parts found at Hazor (Hzr.17387). Notice the concentric splitting following stacking of the growth bands; **D)** another ivory furniture part from Hazor (Hzr.16422). On the left, the distinctive Schreger pattern topography is observed, and on the left, the splitting along straight lines that correspond to the Ra profile (see Fig. 2)

***Step 1: 2*** Observing surface features stemming from the raw material’s unique structure

In bone, the basic identification feature is the holes of the vasculature between bone laminae, which can be seen as a cancellous matrix when the artifacts’ facets correspond to the original bone Tr plane (Fig. 4: C) or as longitudinally cut channels, then we see them in lateral view to the longitudinal bone axis (Fig. 4: A–B, D; Hakker-Orion 1999; Locke 2004).

**Fig. 4.**
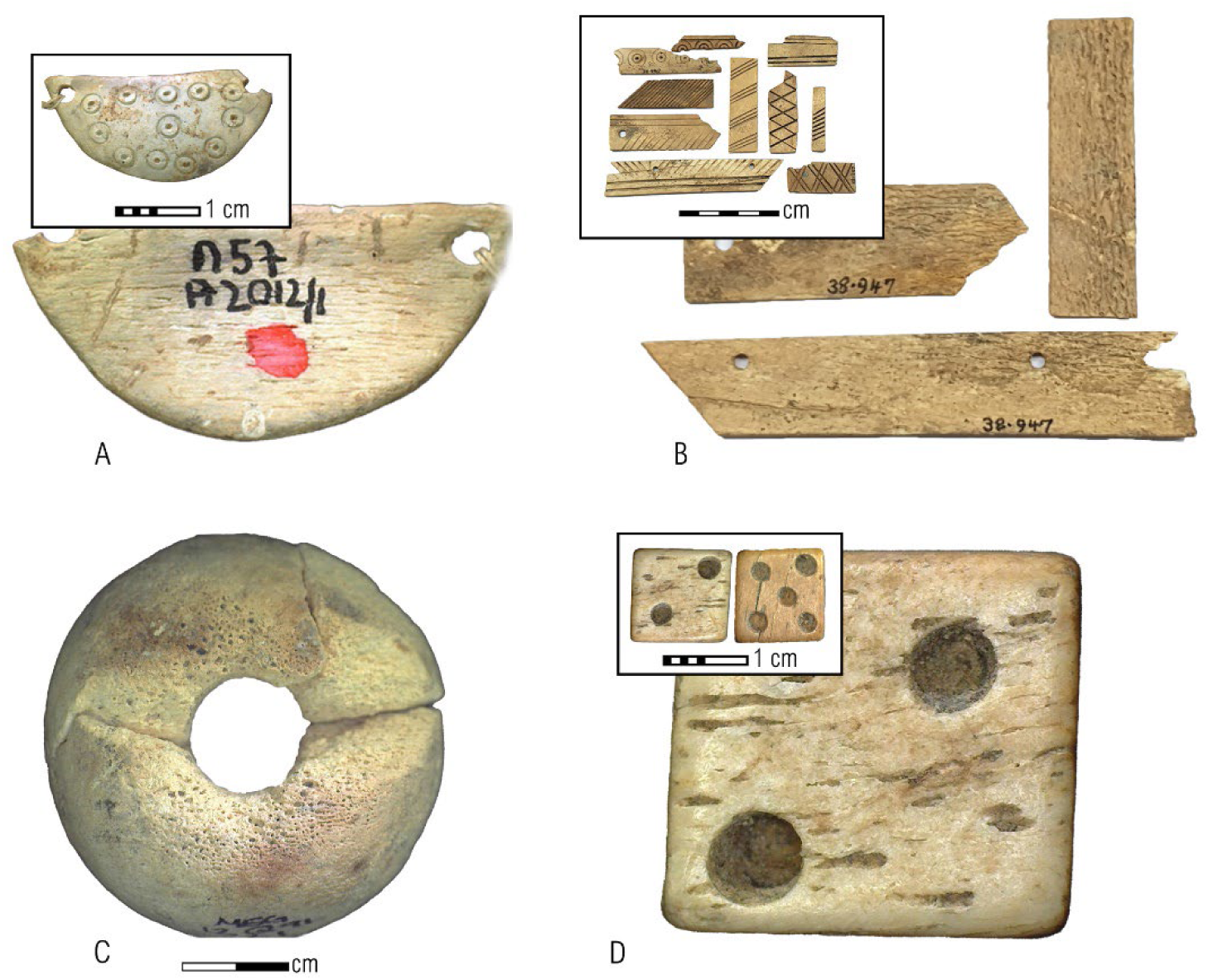
The most straightforward way to identify a bone artifact is the vasculature channels seen on the artifact surface, either as tiny pores or short and parallel carved lines. **A)** A pendant from Iron Age IIB Hazor, Hzr.21-1406; **B)** Furniture decorations from Middle Bronze Age Megiddo, Meg.38-947; **C)** Spindle whorl from Iron Age II Megiddo, Meg.12.Q.091.AR001; **D)** A die from an Iron Age II tomb from Jerusalem, JLM.Hin.80-1259.

In elephant ivory, the composite hierarchical structure resulting in the Schreger columns pattern can sometimes be observed on a macroscopic level (Fig. 5; compare with Baker et al. 2020: figs. 2.6, 2.7; more about this unique pattern, below). The artifact’s facet on which this pattern is identified corresponds to the original tusk Tr plane.

**Fig. 5.**
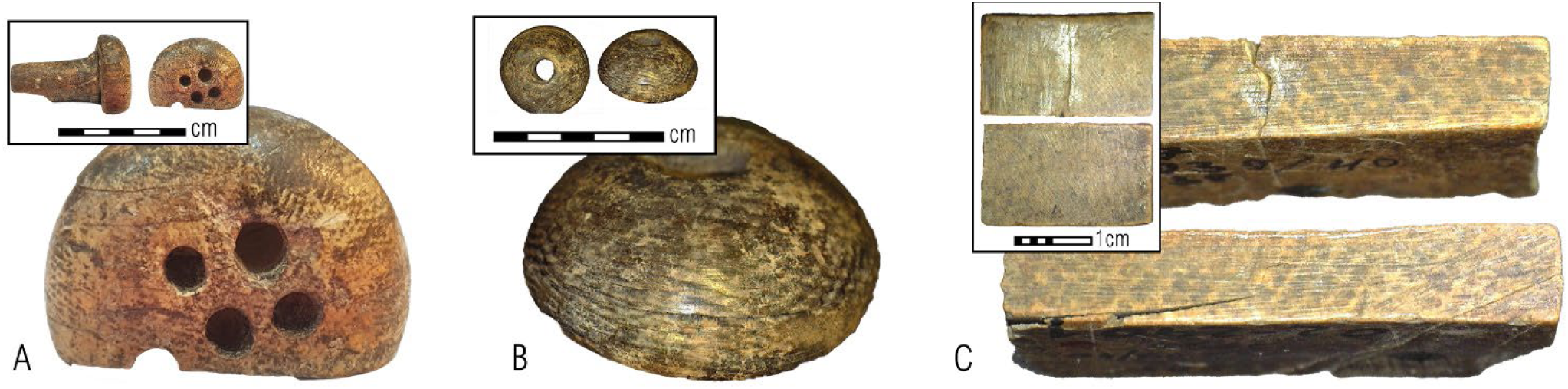
Schreger pattern seen on the surface of selected artifacts. **A)** A possible knob handle from LBA Miqne-Ekron, Mqn.6871; **B)** A spindle whorl from Iron Age II Dor, Dor.96232; **C)** furniture inlays from LBA Lachish, Lcs. 02-2901.

***Step 1: 3*** Observing typical splitting or fracture patterns

Elephant ivory breakage displays a different pattern in each profile sectioning the longitudinal tusk axis. A typical Tr profile breakage can be described as concentric circles following the conical growth bands (Fig. 6). A breakage along the Ra plane is characterized by parallel lines or ‘cone-in-cone’ splitting (Fig. 6: E–F; Krzyszkowska 1990: pl. 5). Identifying these features might also assist with less accessible items, as these fractures can be seen from afar (for instance, behind a museum’s glass display closet) and occasionally enable identification from photos (even older ones).

**Fig. 6.**
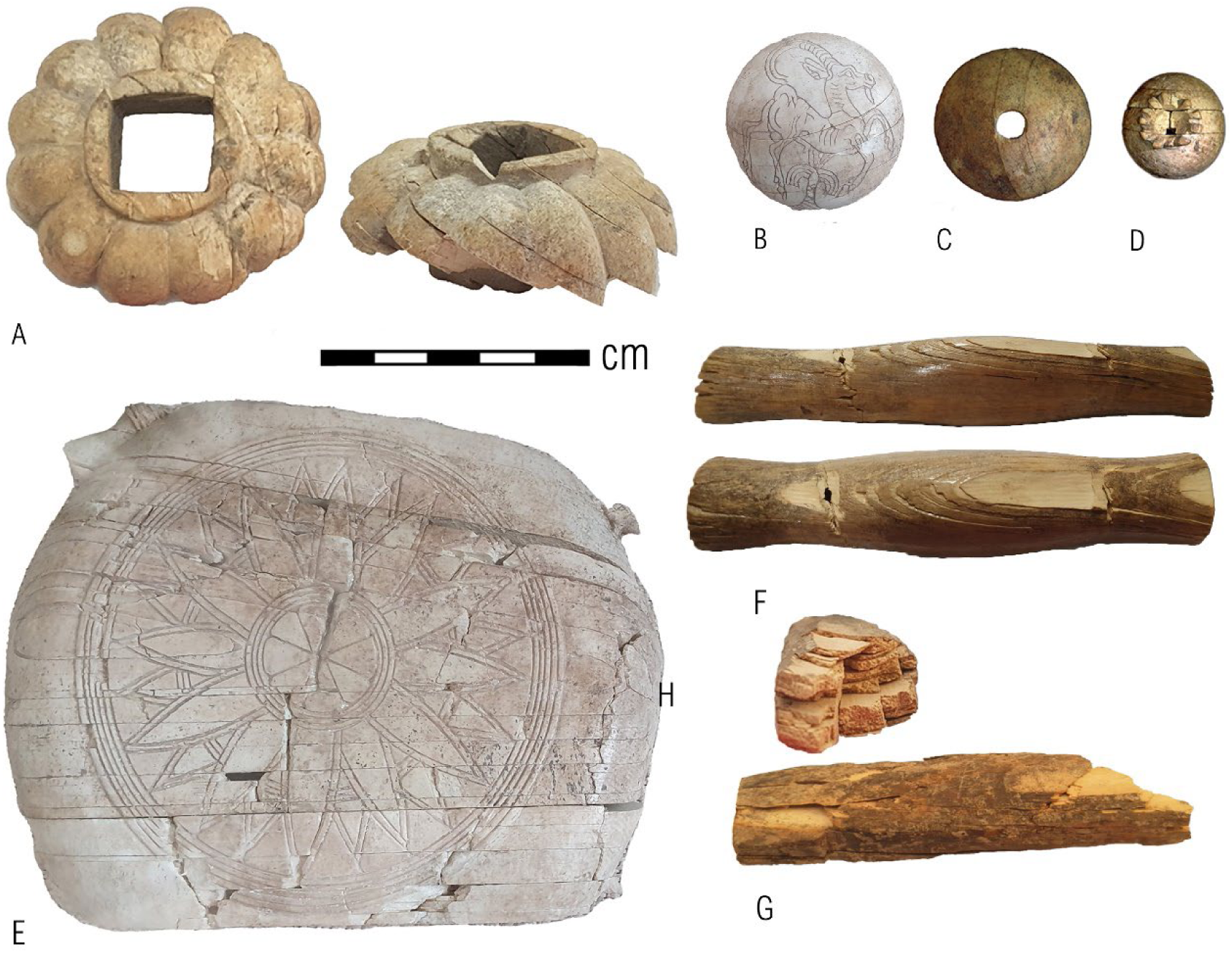
Fracture lines along the growth bands in elephant ivory are seen as curved or concentric lines corresponding to the tusk TR profile and as straight parallel lines on the Ra profile of the original tusk. **A)** Fracture lines on furniture fitting from Iron Age II Keisan seen as curved lines (left) correspond to the Tr plane and seen as straight lines (right) characterizing the Ra plane, Ksn.79-305; **B)** Game piece from LBA Megiddo, Meg.38-792; **C)** Spindle whorl from Iron Age I Dor, Dor.09D2-6331; **D)** Poppy-capsule-shaped finial from Iron age I Askelon, Ask.56588; **E)** The ‘cone-in-cone’ breakage pattern is visible from a distance on an ivory cosmetic container from LBA Megiddo, Meg.38-823 (not to scale). The photograph was taken behind the display closet glass in the Rockefeller Museum, and the lines are also discernable in Loud’s publication from 1939 (pl. 28: 148); **F)** ‘cone-in-cone’ splitting pattern seen on a handle from LBA Beth Shean, Bsn.14-2148; **G)** Fracturing along parallel lines indicates the tusk Ra plane and is seen as curved lines in the artifact cross-section, corresponding to the tusk TR plane. A handle from LBA Miqne-Ekron, Mqn.6576.

The three-dimensional crosshatched breakage pattern is typical of elephant ivory transverse section and is usually best observed using side lighting. This pattern corresponds to the Schreger lines or columns, which is the characteristic hallmark of elephant tusk (Fig. 7). This unique morphology is a result of the intersection of the radially arranged microlaminae in a matrix having cylindrical growth bands (Locke 2008: 430–3).

**Fig. 7.**
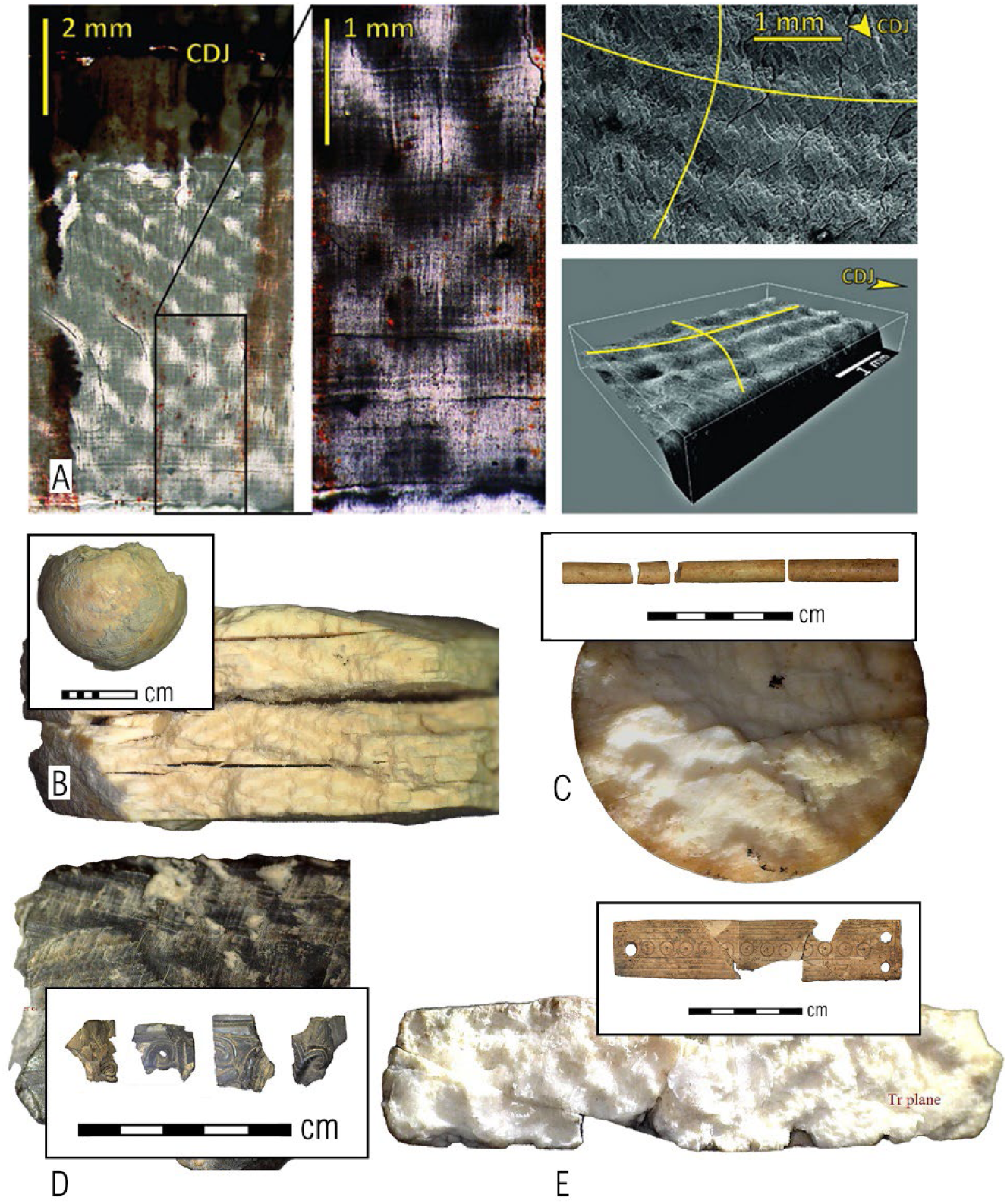
Crosshatched breakage topography corresponding to the Schreger columns in elephant ivory is better viewed using some side lighting and usually on fresh breakages. **A)** SEM visualizations of the Schreger pattern created to explain its natural formation, ‘CDJ’ denotes the cementum-dentine junction (taken from Virág 2012: fig. 3); **B)** A pommel finial from the City of David, Jerusalem, dated to the Iron Age II, CoD.M.973116; **C)** A fractured spindle from Iron Age I Dor, Dor.32299. **D)** A fragment of palmette inlay from Iron Age II Samaria, Smr.12235; **E)** A furniture panel from Iron Age I Dor, Dor.160086.

Artifacts that were carved so they encompass the hippo lower canine complete or partial Tr plane might be fractured or split along a distinctive weakness plane following the teeth Y-shaped ‘forming face’ (Locke 2008: fig. 6). In some publications, the formation face is referred to as the ‘commissure’ (Fig. 8).

**Fig. 8.**
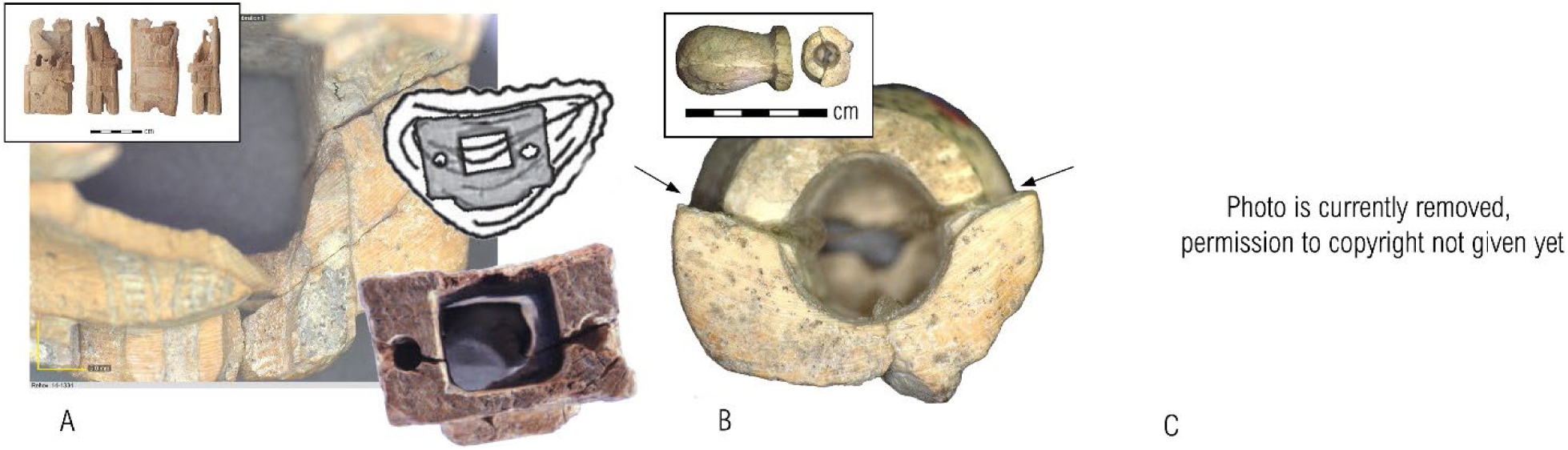
Growth bands angled to the forming face seen on the cross-section of the hippo lower canine (Locke 2008: 427–8). **A)** A fracture created along the forming face of an Iron Age II statuette from Tel Rehov, Rhv.14-1334 (the drawing and below photo adapted from Mazar and Shatil 2020, reproduced with the permission of the Institute of Archaeology, The Hebrew University of Jerusalem); **B)** Similar fracture on a handle from LBA Lachish, Lcs. 05-1626; **C)** Image of the complete lower canine transverse profile, where the angled bands are more clearly seen. The forming face is also referred to as the *Tusk Interstitial Zone*, or TIZ (adapted from Baker et al. 2020: fig. 6.2).

**Step 2**. Microscopic discrimination of ivory from bone

When the compact bone part is used to manufacture artifacts of typological range parallel to ivory, definitive discrimination between bone and ivory based on macroscopic observation may prove unproductive. Polished and worked, the compact bone surface is similar in color and texture to the ivory surface. We can then turn to microscopic inspection, applying a two-stage scrutiny: first to discriminate bone from ivory (*Step 2*), and then to identify the ivory object raw material as elephant or hippo (*Step 3*).

***Step 2: 1*** Targeting the bone micro-morphological indicative features using a magnification resolution of X20–X80.

Two primary features are looked for: the intersecting lines of the vascular bone compartments and the osteons, which comprise the cylinder of concentric lamellae around a tubular space containing blood vessels (Locke 2004).

The observed parallel lines are the vascular bone compartments, which might, at first sight, be viewed as similar to hippo ivory growth bands (see below). However, they attest to the bone identification if transected by Volkmann’s canals, which is observed as perpendicular connecting lines (Fig. 9). This pattern is primarily seen on the radial plane to the long bone longitudinal axis (Locke 2004: fig. 1).

**Fig. 9.**
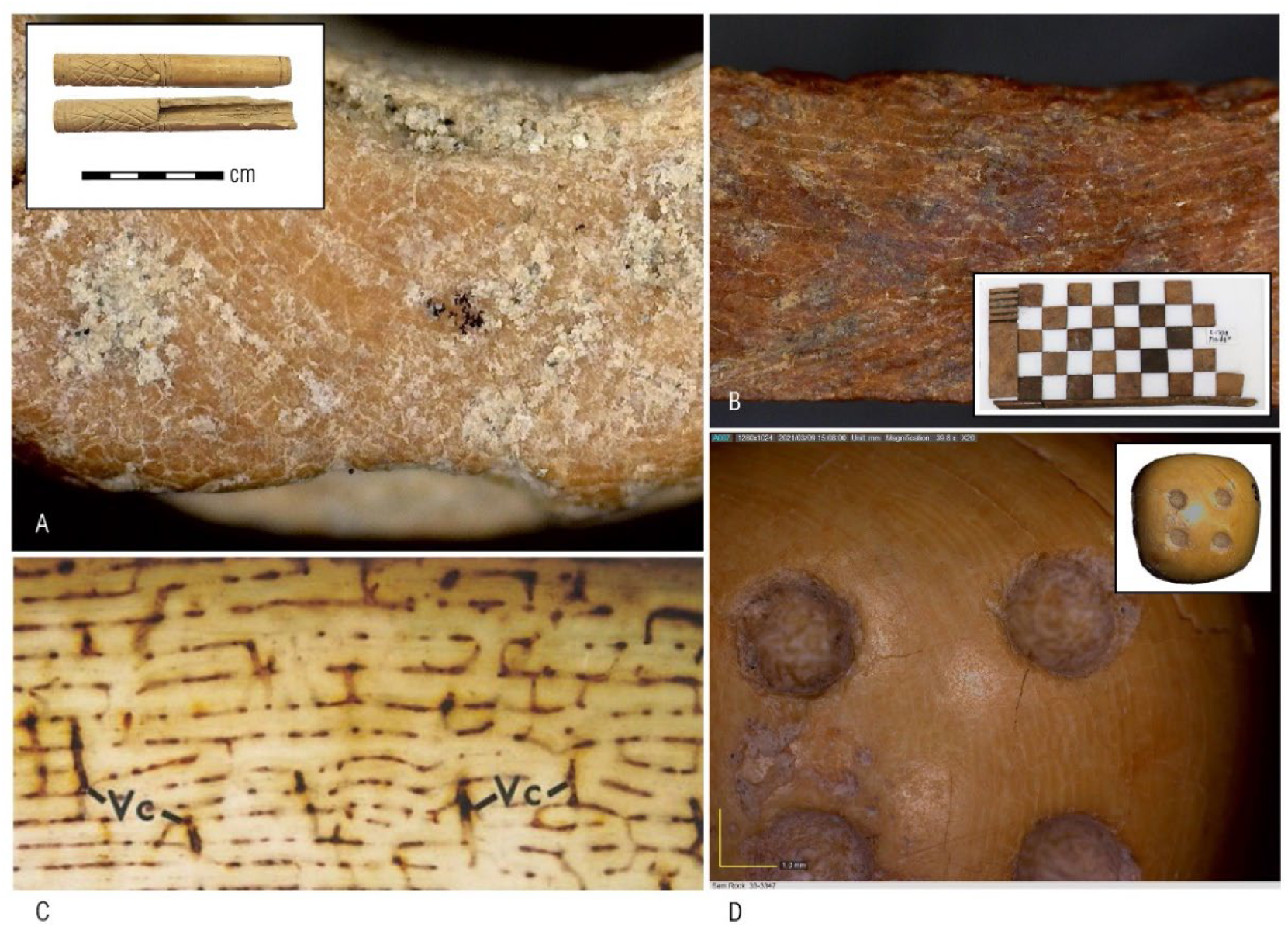
The vascular bone compartment might be seen as parallel (straight or curved) lines, transected by short perpendicular lines, known as the Volkmann’s canals. **A)** Cylindrical handle from Beth Shaen, Bsn.36-1708; **B)** Square inlay from a box from LBA Hazor, Hzr.17368; **C)** Silver-stained in a modern buffalo humerus (adapted from Locke 2004: fig. 1a, ‘Vc’ stands for Volkmann’s canals); **D)** A die from Iron Age II Samaria, Smr.33-3347.

A cellular-shaped arrangement of the osteons (the tiny pores of the vasculature) usually requires magnification higher than X40 (Fig. 10), but when observed, they are a principal feature in discriminating bone from ivory.

**Fig. 10.**
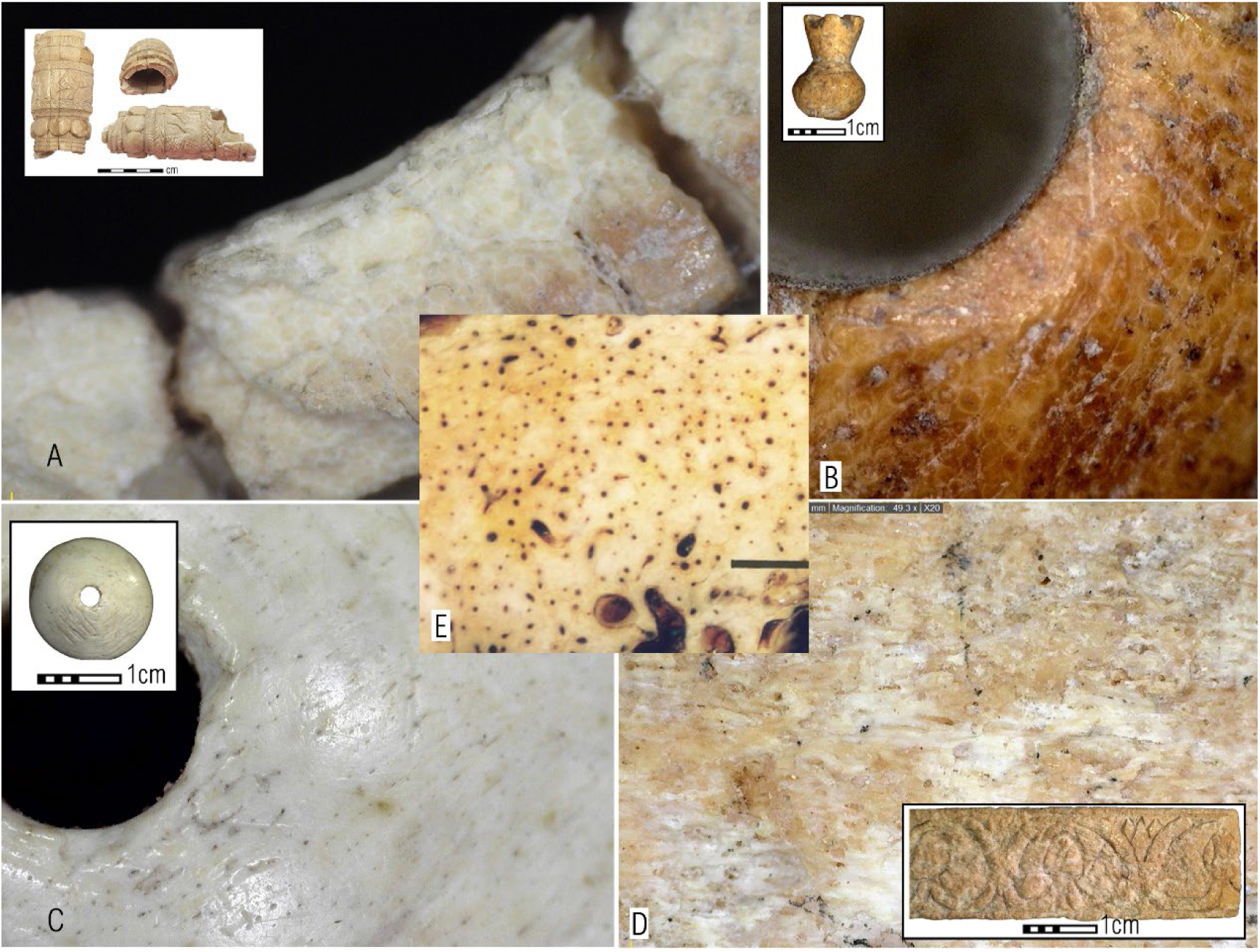
Tiny pores of the osteonic blood vessels (Locke 2004: 4–6); **a)** a handle from Hazor, Hzr.11-1326; **b)** a finial from Miqne-Ekron; **c)** Bead/button from Megiddo, Meg.34-1814; **d)** ivory panel decorated with lotus flower and buds, Ira.99-4378; **e)** Locke 2004: fig. 4A.

**Step 3**. Microscopically identifying elephant and hippo ivory

As mentioned above, ivory is a dentinal tissue, namely a tooth, regardless of its taxonomic classification, and thus might originate from a relatively wide range of species. Nevertheless, as the geographically proximate possible sources for ivories in the Levant are either elephants (African or Asian) and hippos (local or Nilotic), they are the focus here.

***Step 3: 1*** Identifying the Schreger pattern (see above *Step 1: 3*), the most distinctive feature of elephant ivory, sometimes requires higher magnification. The use of side light is recommended to highlight the breakage topography.

***Step 3: 2*** Identifying the ‘feathers’ pattern corresponding to the tusk’s Ta plane.

If the artifact was broken, cracked, or split, so the Ta plane is visible, the ‘feathers’ pattern is a very effective taxonomic discrimination tool. This pattern results from the microlaminae helicoidal arrangement (Locke 2008: 437–41, figs. 13, 15–16), and it is the expected pattern along the splitting planes of the growth bands (Fig. 11). Note that if the artifact was carved closer to the tusk core, or if an oblique plane between the Ta and Ra profiles is exposed (and visible), then instead of ‘feathers,’ the topography has a more wave-like pattern, similar to that of hippo Ta profile (Lock 2008: figs. 14, 16).

**Fig. 11.**
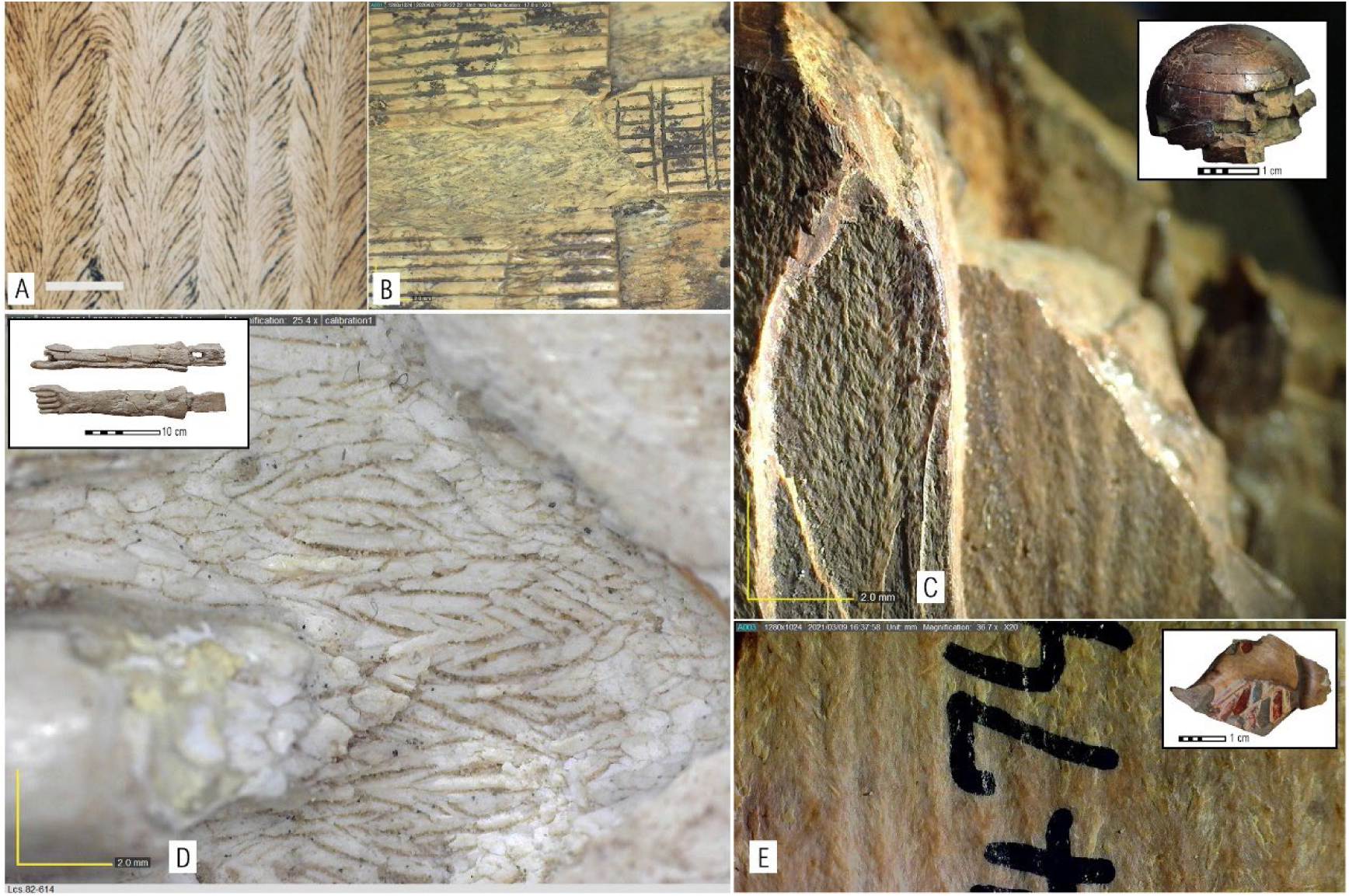
’Feathers’ pattern seen on the artifact facet that corresponds to the elephant tusk Ta plane. **A)** Closer to the tusk core, the stacking of dentinal tubules resembles truncated helicoids, pen marker-stained modern tusk, bar scale 1mm (adapted from Locke 2008: Fig. 15A); **B)** Decorated panel from Middle Bronze Age Hazor, Hzr.28698; **C)** Pommel-shaped finial from the Iron Age II City of David, Jerusalem, CoD.Sh.86-1797; **D)** Modular statuette hand from LBA Lachish, Lcs.82-614; **E)** Colored inlay piece from Iron Age II Samaria, Smr. 35-2474.

Hippo lower canines are more frequently used than the upper incisor in the Southern Levant as a source of ivory. A probable explanation is their larger size compared to the incisors (thus enabling the utilization of more raw material and carving larger objects; Baker et al. 2020: fig. 6.1).

***Step 3: 3*** The most distinctive micro-morphological feature of hippo ivory is the straight growth bands viewed as repeating parallel lines, usually crossing the artifact’s planes (Fig. 12).

**Fig. 12.**
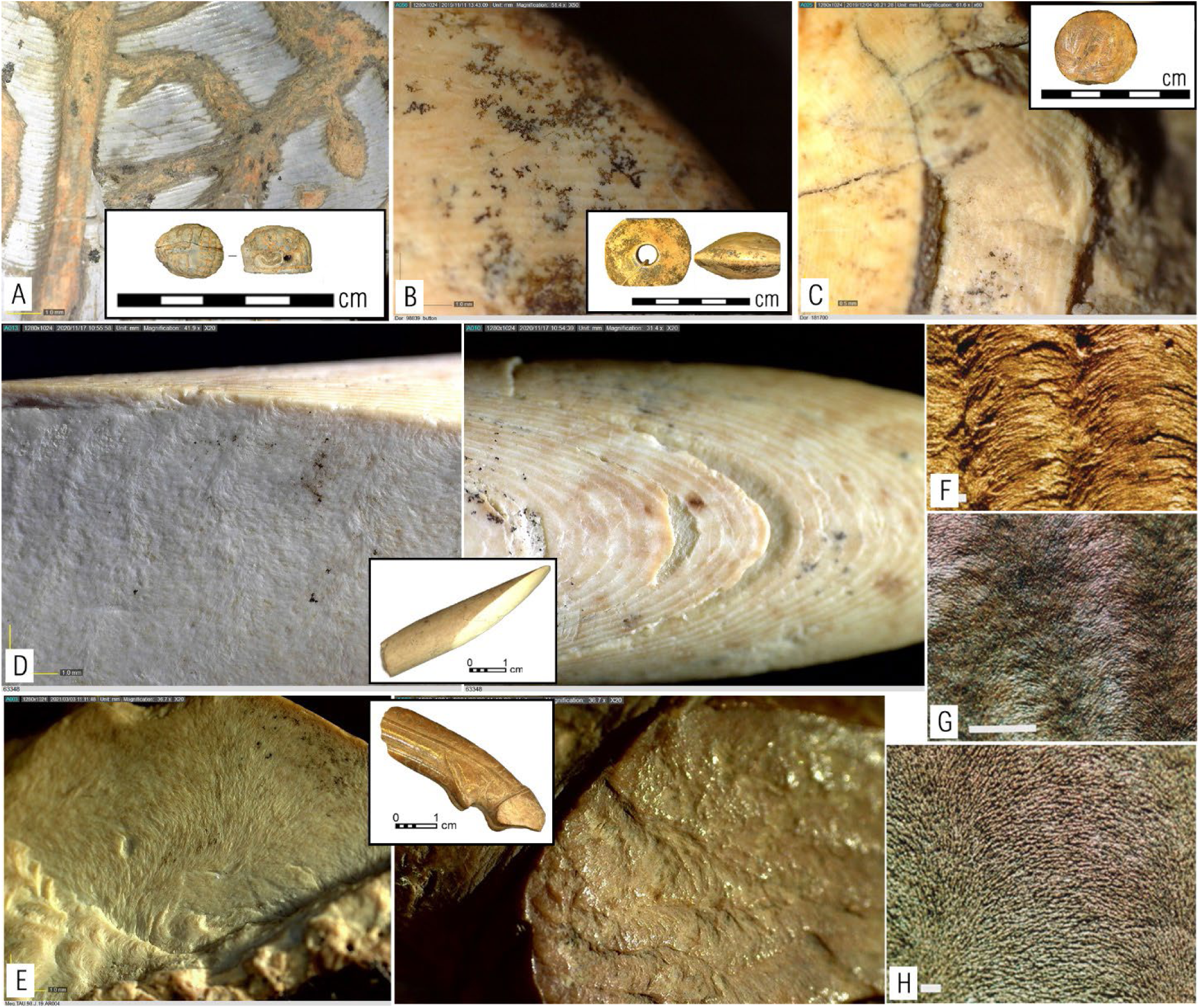
The micro-morphological features that characterize the hippo lower canine are the growth bands viewed as parallel lines and crossing at least two artifact facets (A–C) and the wave-like lines characterizing the breakage pattern of the canine Ta plain. **A)** A seal from Iron Age II Abel Beth-Maacha, Abm.71625; **B)** A bi-conical shaped spindle whorl from Iron Age I Dor, Dor.98839; **C)** Raw material blank made of hippo incisor from Iron Age I Dor, Dor.181700; **D)** Rod fragment from Iron Age I Ashkelon. The parallel growth bands display here a ‘cone-in-cone’ pattern, and the wave-like micro-topography on the artifact cross-section corresponds to the canine Ta plane, Ask.63348; **E)** Wave-like micro-morphology on a fresh break on a possible handle of a cosmetic spoon from Middle Bronze Age II Megiddo, Meg.98.J.019.AR004; F–H) The surface irregularity of the micro-topography of the dentinal tubules stacking in Ta plane, modern hippo lower canine stained by marker pen (Locke 2008: fig. 8A–C).

The growth bands correlate with the original teeth Tr profile (Fig. 12: A–B; Locke 2008: 427–30, fig. 6), and sometimes the Y-shaped forming face can also be viewed (see above, Fig. 8).

***Step 3: 4*** Searching for the hippo lower canine wave-like micro-topography seen on the Ta profile, resulting from the dentinal tubules traversing the dentine matrix in a helicoidal arrangement (Fig. 12: D–E; Locke 2008: figs. 8, 9).

***Step 3: 5*** The Hippo incisor is less common, and the observed micro-morphological feature is the concentric growth bands (Fig. 12: C; Locke 2008: 426–7; Baker et al. 2020: fig. 6.3).

Some artifacts’ preservation state enabled the identification of some or all the micro-morphological attributes, thus enabling a three-dimensional perspective to correlate the artifacts with the original tusk sectioning profiles and validate our observations. Such an outlook is summarized in Fig. 13. An illustrated scheme of the three-step identification is provided in Fig. 14.

**Fig. 13.**
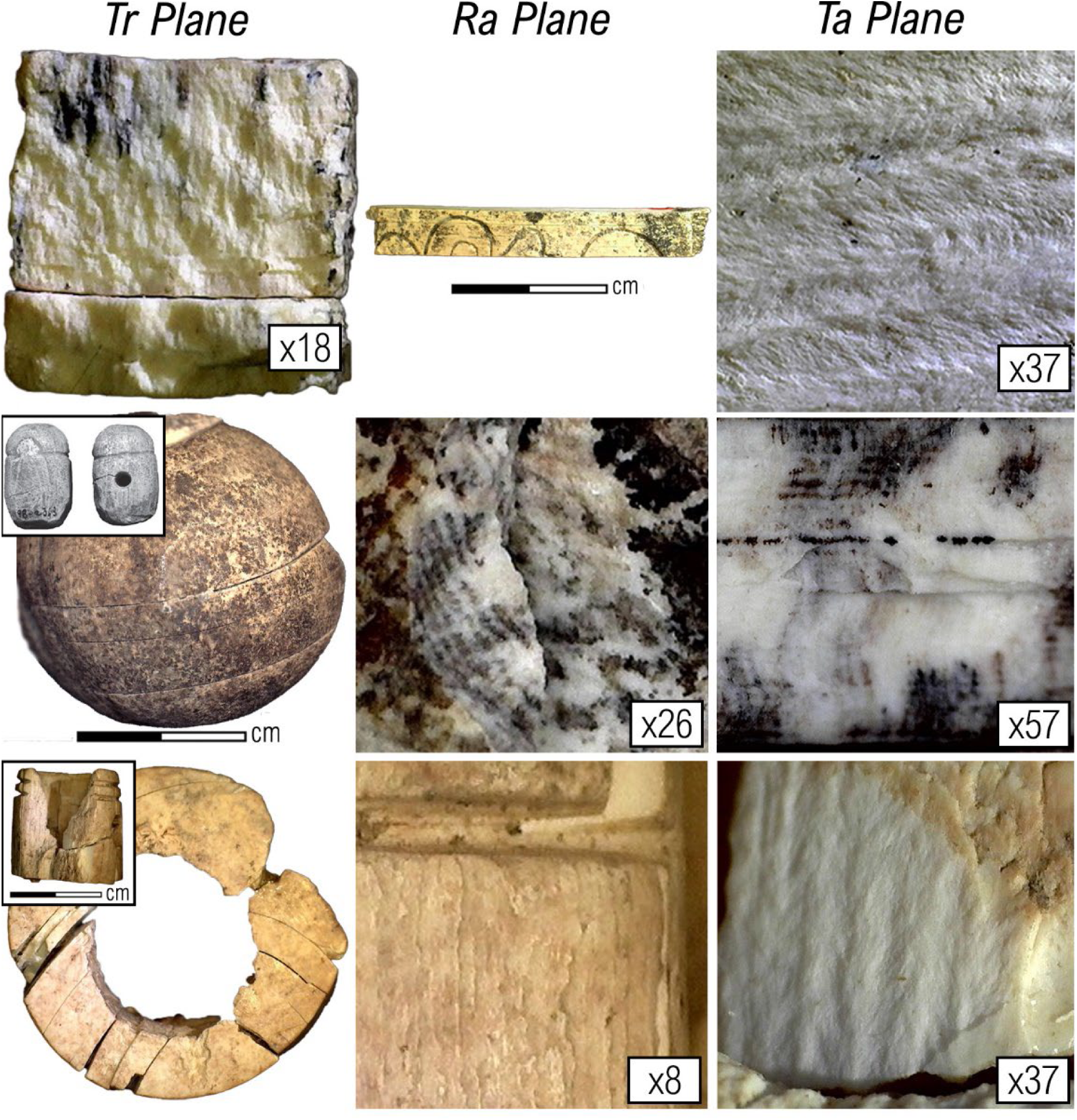
Conclusive outlook of the different micro-morphological features characterizing each of the original elephant tusk profiles. The magnification used for each artifact’s facets is provided and can be considered a rough average needed to identify the requested attributes. **Upper row:** An inlay from Iron Age I Ashkelon, Ask.56705. Note: The Ta plane wave-like pattern indicates the artifact was carved closer to the tusk core. **Middle row:** A fitting from Iron Age II Dor, Dor.93986. The Schreger pattern is visible to the naked eye, indicating the Tr plane of the tusk. The Ta plane brakeage shows the wave-like arrangement of the dentinal tubules stacking. **Lower row:** A fitting from Iron Age I Ashkelon, Ask.48456. The ‘feather’ pattern observed on the Ta plane indicates the artifact was carved closer to the tusk periphery. The growth band angles on the Tr plane indicate an oblique angle to the tusk cross-section.

**Fig. 14.**
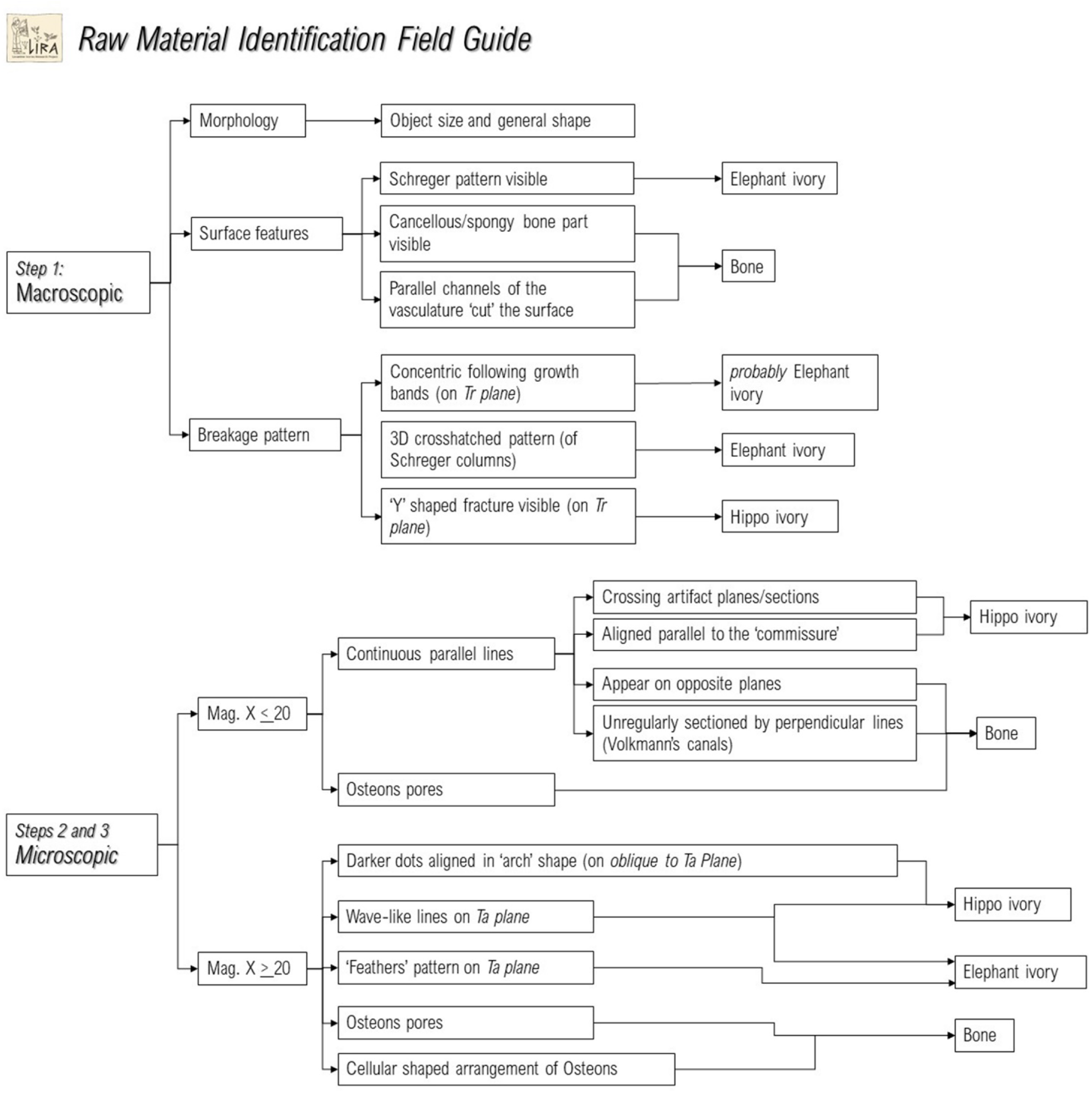
An illustrated scheme of the identification protocol.

## ZooMS analysis authenticates the microscopic identification

Ivory artifacts are rare finds within the material culture assemblage, and when found intact or in restorable shape, they are in high demand for display in museums, and even when not, they are usually well curated. Therefore, destructive sampling of ivory artifacts is bound to strict limitations and follows the basic instruction: not to harm intact artifacts; when sampled, to acquire the minimum needed material from the minimum amount of objects; and, when possible, to take micro-fragments that already departed from the original artifact due to their preservation state (relevant only for fragments unequivocally proven to belong to the original artifact, for example, stored in a container separated from other artifacts).

The proteomic identification for 51 ivory artifacts out of the total 58 successfully identified samples matched that of the microscopic identification, and two more artifacts were correctly identified as bone. Together, they consist of 91% of correct identifications and attest to the integrity and validity of the microscopic method. The proteomic analysis eventually proved only five microscopic identifications erroneous (Table 1): two elephant ivory inlays were erroneously identified as bone; one elephant ivory rod was identified as hippo; one bone rod was identified as elephant ivory, and one boar canine was identified as hippo (see Shochat 2023: 291–295 for an elaborated introduction to this method, and pp. 361–364, App. A for the analysis results).

**Table 1.** The integrity of the microscopic taxonomic identification, as corroborated by the ZooMS analysis.

| **ZooMS vs. microscopic identification** | **N** | **%** |
| --- | --- | --- |
| Successfully identified by ZooMS | 58 |  |
| Ivory identification **match** microscopic | 51 | 88% |
| Ivory identification **unmatch** microscopic | 5 | 9% |
| Correctly identified as bone | 2 | 3% |

## Summary

For over a century, scholars advocated ivory’s status as a highly valued material, a charismatic medium broadcasting rulership and authority, and associated with society’s elite strata. Its prestige probably stemmed from its dependency on control over sources of power and wealth to obtain it (Aubet 2013: 94–6), usually from afar. Despite the cost and technological challenges of far-flung trade, ivory was exchanged owing to its cultural significance and contribution to social structuring. Scrutinizing ivory artifacts distribution trends – taxonomic, chronological, stratigraphic, and contextual – and tracing their geographic origins thus bear great potential for illuminating intercultural exchange-related aspects of past societies.

The method described here provides a straightforward, non-destructive, and widely applicable practice to examine large-scale assemblages of archaeological ivories. Furthermore, until bio-molecular analyses, namely proteomic, stable isotopes, and aDNA, become more accessible for a broader range of archaeologists, it can be effectively used as a prescreening method.

## Acknowledgments

I would like to acknowledge a great number of scholars and fellow researchers for their kind reception and generosity in giving me access to examine the materials in their keeping: first and foremost, Miki Sebanne, Alegre Savariago and Debi Ben-Ami of the Israel Antiquity Authority National Treasure Department; Ilan Sharon and Sveta Matskevich of the Hebrew University of Jerusalem and the Tel Dor Project; Bracha Zilberstein of the Mizgaga Museum at Nahsholim; Dafna Tzoran and Mimi Lavi of the Hebrew University; Shlomit Bechar and Amnon Ben-Tor of the Hebrew University Renewed Expedition to Hazor; Shira Albaz and Aren Maeir, Bar Ilan University, of the Tell es-Safi/Gath Archaeological Project; Assaf Kleiman and Yana Kirilov of the Tel Aviv University Megiddo Expedition; Ido Koch and Ze’ev Herzog, Tel Aviv University, for access to the Tel Arad material; Liora Freud, Sabina Kleiman, Oded Lipschits, and Alexandra Wrathall, Tel Aviv University, for access to Tel Azekah and Ramat Rahel material; Sy Gitin and Anna de Vincenz for access to Tel Miqne/Ekron assemblage; Levana Zias, David Ilan, Yifat Thereani and Rahel Ben-Dov of the Hebrew Union College for access to the Tel Dan material; Michal Artzy, University of Haifa, for the access to the Tel Nami collection; Golan Shalvi, Ben-Gurion University for the Tel Shiqmona materials; Nitza Bashkin Joseph, curator of the Eretz Israel Museum, for access to the Tel Qasile and Timna materials; Nurit Goshen of the Israel Museum, for the opportunity to examine the collections from Lachish; and Yiftah Shalev, Israel Antiquity Authority, and Yuval Gadot, Tel Aviv University, for the access to the wonderful ivories from Givati parking lot, Jerusalem.

## Footnotes

1 The following description is accompanied by photos taken by the author during research on Southern Levantine ivory exchange during the Late Bronze and Iron Age (for more details on the specific items, see Shochat 2023).

